# Stabilization and ATP-binding for tandem RRM domains of ALS-causing TDP-43 and hnRNPA1

**DOI:** 10.1101/2020.12.21.423780

**Authors:** Mei Dang, Yifan Li, Jianxing Song

## Abstract

TDP-43 and hnRNPA1 contain tandemly-tethered RRM domains, which not only functionally bind an array of nucleic acids, but also participate in aggregation/fibrillation, a pathological hallmark of various human diseases including ALS, FTD, AD and MSP. Here, by DSF, NMR and MD simulations we systematically characterized stability, ATP-binding and conformational dynamics of TDP-43 and hnRNPA1 RRM domains in both tethered and isolated forms. The results reveal three key findings: 1) very unexpectedly, upon tethering TDP-43 RRM domains become dramatically coupled and destabilized with Tm reduced to only 49 °C. 2) ATP specifically binds TDP-43 and hnRNPA1 RRM domains, in which ATP occupies the similar pockets within the conserved nucleic-acid-binding surfaces, with the affinity higher to the tethered than isolated forms. 3) MD simulations indicate that the tethered RRM domains of TDP-43 and hnRNPA1 have higher conformational dynamics than the isolated forms. Two RRM domains become coupled as shown by NMR characterization and analysis of inter-domain correlation motions. The study explains the long-standing puzzle that the tethered TDP-43 RRM1-RRM2 is particularly prone to aggregation/fibrillation, and underscores the general role of ATP in inhibiting aggregation/fibrillation of RRM-containing proteins. The results also rationalize the observation that the risk of aggregation-causing diseases increases with aging.

---

Aggregation/fibrillation of TDP-43 in the cytoplasm of neurons is a pathological hallmark of ∼97% amyotrophic lateral sclerosis (ALS) and ∼45% frontotemporal dementia-TDP (FTLD-TDP), that lack any efficacious medicine so far (1-3). TDP-43 is a well-known member of heterogeneous nuclear ribonucleoproteins (hnRNP) containing the folded RNA-recognition motif (RRM), which constitutes one of the most abundant domains in eukaryotes (3-5). Interestingly, ∼70 human RRM-containing proteins including FUS, TDP-43 and hnRNPA1 additionally have the intrinsically disordered prion-like domains of low-complexity sequences with amino acid compositions similar to those of the prion domains in yeast responsible for driving the formation of infectious conformers (2-4). Remarkably, these proteins not only functionally mediate direct interactions with various nucleic acids to control both RNA processing and gene expression, but their aggregation/fibrillation is pathologically characteristic of an increasing spectrum of human diseases including amyotrophic lateral sclerosis (ALS), frontotemporal dementia (FTD), Alzheimer’s disease (AD), chronic traumatic encephalopathy, muscle regeneration/degeneration and multisystem proteinopathy (MSP) (1-10).

414-residue TDP-43 contains two folded RRM domains (Fig. 1A), which are not only essential for recognizing UG-rich sequences near RNA splice sites (11), but also extensively demonstrated to participate in disease-causing aggregation/fibrillation in additional to its C-terminal prion-like domain (11-19). Remarkably, small molecules targeting the RRM domains of TDP-43 have been recently identified to reduce locomotor defects in drosophila model of ALS (19). We also found that ATP, the universal energy currency, specifically binds the RRM domains of FUS and TDP-43 to inhibit their amyloid fibrillation (20-22). Therefore to understand the factors and mechanisms governing their aggregation/fibrillation is not only of fundamental interest, but represents an essential step for further development of therapeutic strategies/molecules to treat these diseases.

**Fig. 1.**
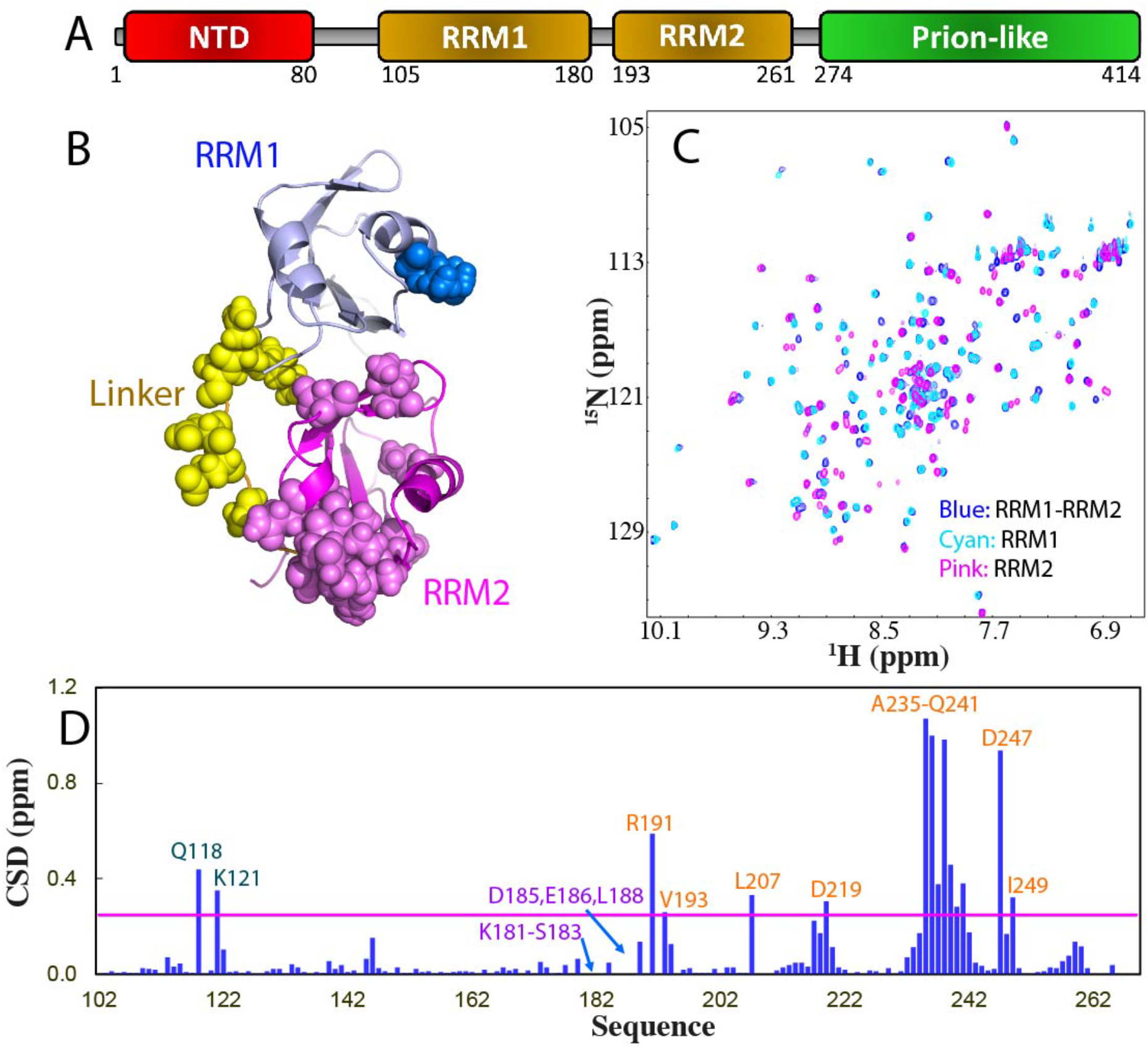
Dissection-induced perturbation of TDP-43 RRM domains. (A) 414-residue TDP-43 contains: N-terminal domain (NTD) over residues 1-80; two tandemly-tethered RNA recognition motifs (RRM1 and RRM2) over residues 102-269, and C-terminal prion-like domain over residues 274-414. (B) NMR structure (PDB ID of 4BS2) of the TDP-43 RRM1 (light blue) and RRM2 (pink) structure in which the residues are displayed in sphere for those with HSQC peaks disappeared (yellow), significantly shifted in RRM1 (blue) or in RRM2 (light pink) upon dissection. (C) Superimposition of ^1^H-^15^N NMR HSQC spectra of the ^15^N-labeled tethered RRM1-RRM2 domains (blue), the isolated RRM1 (cyan) and RRM2 (pink) proteins at 50 μM in 10 mM sodium phosphate buffer (pH 6.8) containing 150 mM NaCl and 10 mM DTT. (D) Residue-specific chemical shift difference (CSD) of the RRM1 and RRM2 domains between the tethered and isolated forms. Significantly shifted residues are labeled and displayed as spheres in (B), which are defined as those with the CSD values > 0.25 (average value + one standard deviation) (pink line).

Structurally, two TDP-43 RRM domains adopt the same overall fold shared by all RRM domains, which consists of a four-stranded β-sheet and two perpendicular α-helices (Fig. 1B). Recently, we showed that aggregation/fibrillation of the single FUS RRM domain not only depends on thermodynamic stability, but also on conformational dynamics (23). Briefly, the FUS RRM domain undergoes fibrillation not only due to its relatively low stability with melting temperature (Tm) of 55 °C, lower than that of the human *γ*S-crystallin with Tm of 71 °C (24), but also resulting from its high conformational dynamics, that allow the dynamic opening of the structure to expose its central hydrophobic regions for aggregation/fibrillation (23). Consequently, although ATP induces no enhancement of the thermal stability of the FUS RRM domain, it is sufficient to kinetically inhibit the amyloid fibrillation through weak but specific binding with a Kd of 3.77 mM to a pocket within the conserved nucleic-acid-binding surfaces, which thus leads to blocking the dynamic opening of the structure (19).

It has been a long-standing puzzle that the tethered RRM domains of TDP-43 were particularly prone to aggregation (11) while the isolated RRM1 and RRM2 domains appeared to be stable and soluble as evidenced by their NMR structures determination at high protein concentrations by RIKEN Structural Genomics/Proteomics Initiative (PDB ID of 2CQG and 1WFO). We also observed that in our buffer mimicking *in vivo* conditions (10 mM sodium phosphate at pH 6.8 with 150 mM NaCl and 10 mM DTT), the tethered TDP-43 RRM1-RRM2 could form amyloid fibrils within several days while the isolated RRM1 and RRM2 failed even after a month (22). In particular, we found that under exactly the same conditions, the tethered RRM1-RRM2 of TDP-43 has only one unfolding transition with Tm of only 49 °C, lower than Tm of the isolated RRM1 (57 °C). On the other hand, the binding affinity of ATP has been characterized to be higher to the tethered RRM1 than the isolated RRM1 (21,22).

So far, it remains completely unexplored for the relationship of thermodynamic stability, conformational dynamics and ATP-binding of RRM domains between the tethered and isolated forms. In the present study, with experimental methods including DSF and NMR as well as molecular dynamics (MD) simulations, we aimed to address this problem by characterizing the thermal stability, ATP-binding and conformational dynamics of TDP-43 and hnRNPA1 RRM domains in both tethered and isolated forms. The most unexpected finding is that upon tethering, TDP-43 RRM1 and RRM2 become highly coupled to behaving as one unfolding unit with the stability significantly reduced. By contrast, no significant destabilization was observed upon tethering of two RRM domains of hnRNPA1. Our study further showed that the tethering-induced effects mainly result from the inter-domain interactions as detected by NMR characterization and analysis of inter-domain correlation motions calculated from MD simulations. Moreover, for the first time, we found that ATP can also specifically bind the hnRNPA1 RRM2 but not RRM1 domain with the affinity and complex structure highly similar to those for FUS and TDP-43 RRM domains. Intriguingly, the previous and present results together revealed that ATP bind the RRM domains of both TDP-43 and hnRNPA1 with the affinity higher for the tethered than for the isolated ones. Our study provides the first mechanistic insight into the tethering-induced effects on the tandem RRM domains, and highlights the general role of ATP in inhibiting aggregation/fibrillation of RRM-containing proteins, which extensively causes various human diseases by “loss of functions” or/and “gain of function”.

## Results

### Dissection-induced perturbation of TDP-43 RRM1 and RRM2 domains

As shown in Fig. 1A and 1B, TDP-43 contains two RRM1 and RRM2 domains respectively over residues 105-180 and 193-261 connected by an unstructured linker over residues 181-192. Comparison of NMR structures determined in the tethered (PDB ID of 4BS2) and isolated forms (2CQG for RRM1 and 1WFO for RRM2) revealed that RRM1 has Cα atom RMSD value of 1.63 Å but RRM2 only 0.82 Å, indicating that the overall structures of both RRM1 and RRM2 domains of TDP-43 are well-folded and adopt the same RRM fold in both forms.

Here we cloned and expressed the tethered RRM1-RRM2 of TDP-43 (102-269), as well as its isolated RRM1 (102-191) and RRM2 (191-269). Indeed as extensively observed, the tethered RRM1-RRM2 protein was prone to aggregation even at concentrations of ∼100 μM but the isolated RRM1 and RRM2 proteins showed no significant aggregation at concentrations of ∼1 mM. Nevertheless, the tethered RRM1-RRM2 protein has a well-dispersed HSQC spectrum typical of a well-folded protein at 50 μM in 10 mM sodium phosphate buffer containing 10 mM DTT and 150 mM NaCl (pH 6.8) which mimics *in vivo* environments (Fig. 1C). Two isolated RRM1 and RRM2 proteins also have well-dispersed HSQC spectra at the same protein concentration in the same buffer. However, upon superimposing three HSQC spectra, some significant changes were identified (Fig. 1C): 1) HSQC peaks of residues K181-S183, D185, E186 and L188 over the linker, which were undetectable in the tethered form became detectable in the isolated form; 2) upon dissection some HSQC peaks were largely shifted and the residues with significant chemical shift difference (CSD) are mainly located within the RRM2 domain (Fig. 1D and 1B). This observation strongly suggests that in the tethered form, two RRM domains of TDP-43 have dynamic inter-domain interactions. Consequently although the dissection resulted in no disruption of the overall RRM fold for both RRM1 and RRM2, it did lead to the changes of local conformations, or/and dynamics or/and chemical environments of a set of RRM residues, thus leading to significant shifts of their HSQC peaks.

### Thermal stability and ATP-binding of the isolated TDP-43 RRM2 domain

Previously we have characterized the thermal stability and ATP binding of the tethered RRM1-RRM2 (21) and isolated RRM1 (22) of TDP-43. Intriguingly, the tethered RRM1-RRM2 has only one thermal unfolding transition with Tm of 49 °C, which increased to 54 °C with addition of ATP. By contrast, the isolated RRM1 also has only one thermal unfolding transition but with Tm of 57 °C, which increased to 60 °C with addition of ATP. To understand this unexpected observation, here we measured the thermal stability of the isolated RRM2 under the same conditions. Interestingly, the isolated RRM2 has also only one thermal unfolding transition with Tm of 59 °C, which is not affected by the addition of ATP even up to 15 mM (Fig. 2A). This result suggests that upon tethering, the RRM1 and RRM2 domains of TDP-43 became significantly coupled to acting as one unfolding unit as well as thermodynamically destabilized.

**Fig. 2.**
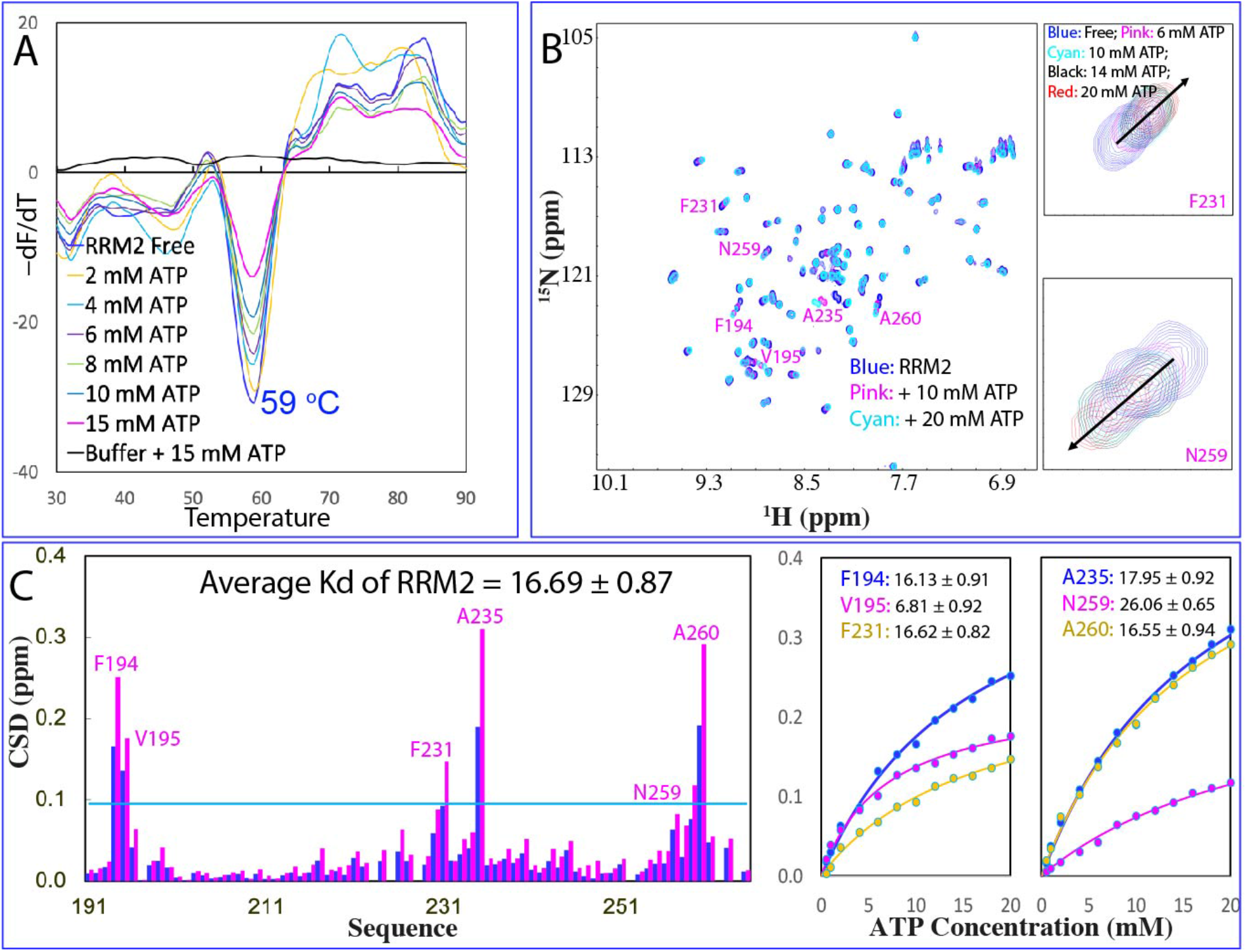
Thermal stability and ATP binding of the isolated TDP-43 RRM2. (A) DSF melting curves of thermal unfolding of the isolated RRM2 domain in the presence of ATP at different concentrations. (B) ^1^H-^15^N NMR HSQC spectra of the ^15^N-labeled RRM2 domain at 50 μM (blue) in 10 mM sodium phosphate buffer (pH 6.8) containing 150 mM NaCl and 10 mM DTT in the absence (blue), and in the presence of ATP at 10 (pink) and 20 (cyan) mM. Traces of two representative HSQC peaks, which have significant shifts but are not severely overlapped with other peaks. For clarity, only peaks at five ATP concentrations are shown: in the free state (blue); in the presence of ATP at 6 mM (pink); 10 mM (cyan); 14 mM (black); and 20 mM (red). (C) Residue-specific chemical shift difference (CSD) of the isolated RRM2 in the presence of ATP at 10 mM (blue) and 20 mM (pink). Significantly shifted residues are labeled, which are defined as those with the CSD values at 20 mM ATP > 0.1 (average value + one standard deviation) (cyan line). Fitting of the residue-specific dissociation constant (Kd): experimental (dots) and fitted (lines) values for the chemical shift differences induced by addition of ATP at 0.5, 1.0, 2.0, 4.0, 6.0, 8.0, 10.0, 12.0, 14.0, 16.0, 18.0 and 20.0 mM.

On the other hand, in the tethered form, the Kd value of RRM1 binding to ATP is 2.58 mM (21) while the Kd of the isolated RRM1 is 3.89 mM (22). Although in the tethered form, RRM2 also has an ATP-binding pocket but with much lower affinity than RRM1, with Kd of 13.85 mM. Here our NMR titrations showed that isolated RRM2 is also able to bind ATP to induce large shifts of a set of HSQC peaks (Fig. 2B) with the overall pattern of the perturbed residues highly similar to that of in the tethered form but with slightly larger Kd (16.69 mM) (Fig. 2C). It is worthwhile to note that ATP binds to both RRM1 and RRM2 domains of TDP-43 with the higher affinity in the tethered form than those in the isolated form.

### Dissection-induced perturbation of hnRNPA1 RRM1 and RRM2 domains

To understand whether the destabilization observed on the tethered TDP-43 RRM1 and RRM2 domains is unique to TDP-43 or also applicable to other RRM-containing proteins. Here we decided to further characterize the tethered and isolated tandem RRM domains of 320-residue hnRNPA1, which contains two RRM domains over residues 15-90, 106-179 respectively (Fig. 3A). Previously, the NMR structures of linked RRM1-RRM2 (25) as well as isolated RRM1 and RRM2 domains (26) have been determined by NMR. The RRM1 domain has Cα atom RMSD value of only 0.68 Å while the RRM2 domain has RMSD of 0.62 Å between the tethered and isolated forms. Interestingly, the linker for the hnRNPA1 RRM1 and RRM2 domains is not completely unstructured as that of TDP-43 but has a short helix over residues 91-96 (Fig. 3B).

**Fig. 3.**
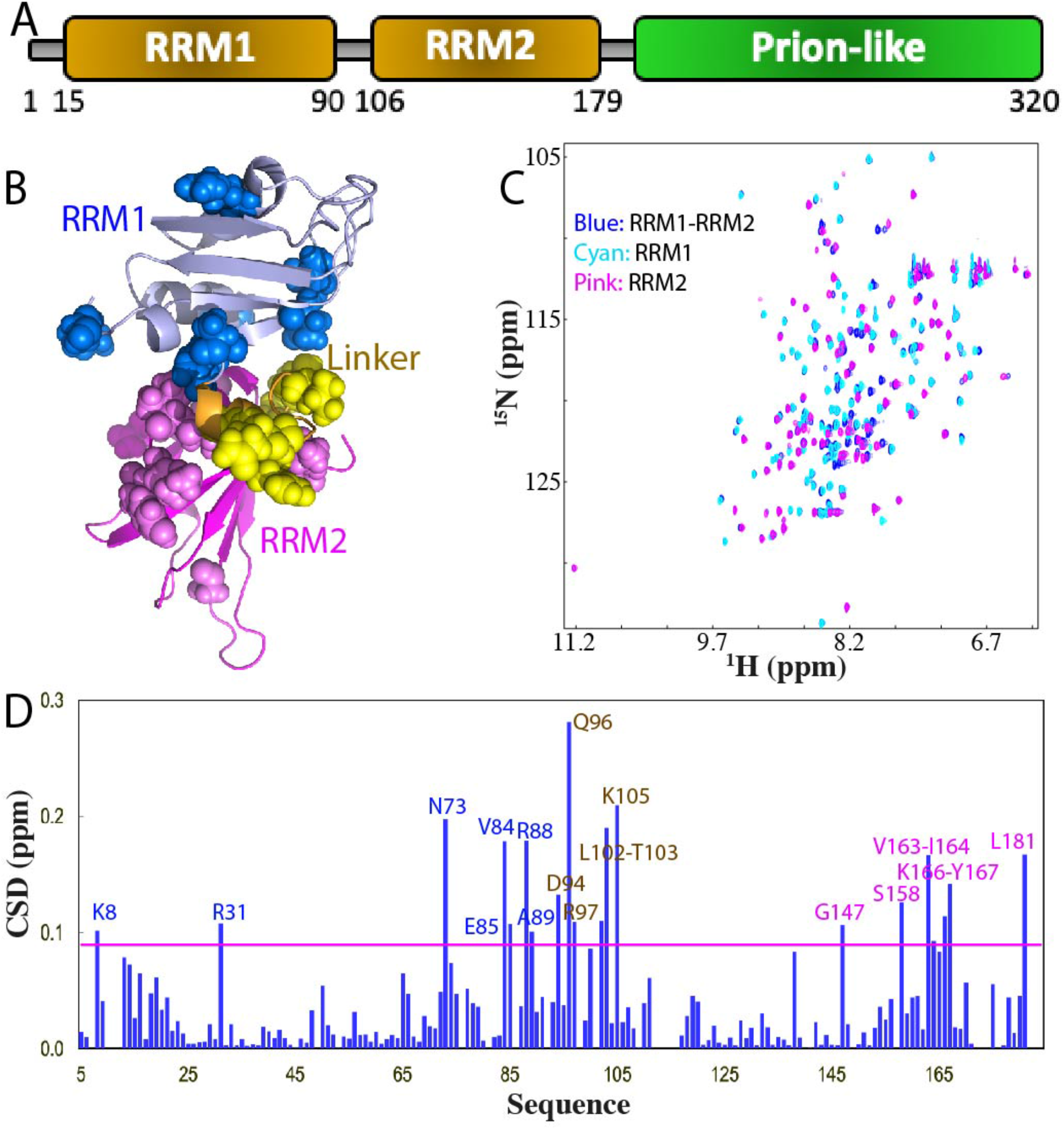
Dissection-induced perturbation of hnRNPA1 RRM domains. (A) 320-residue hnRNPA1 contains: two RNA recognition motifs (RRM1 and RRM2) over residues 1-179, and C-terminal prion-like domain over residues 180-320. (B) NMR structure (PDB ID of 2LYV) of the hnRNPA1 RRM1 (light blue) and RRM2 (pink) in which the residues are displayed in spheres for those with HSQC peaks significantly shifted in RRM1 (blue) or in RRM2 (light pink) upon dissection. (C) Superimposition of ^1^H-^15^N NMR HSQC spectra of the ^15^N-labeled tethered RRM1-RRM2 domains (blue), the isolated RRM1 (cyan) and RRM2 (pink) domains at 50 μM in 10 mM sodium phosphate buffer (pH 6.8) containing 150 mM NaCl and 10 mM DTT. (D) Residue-specific chemical shift difference (CSD) of the RRM1 and RRM2 domains between the tethered and isolated forms. Significantly shifted residues are labeled and displayed as spheres, which are defined as those with the CSD values > 0.09 (average value + one standard deviation) (pink line).

Here we cloned and expressed the tethered hnRNPA1 RRM1-RRM2 (5-184), as well as its isolated RRM1 (5-95) and RRM2 (94-184). Different from what was observed on the TDP-43 RRM domains, the tethered RRM1-RRM2 is highly soluble and has a well-dispersed HSQC spectrum typical of a well-folded protein at 50 μM (Fig. 3C), while two isolated RRM1 and RRM2 proteins are also highly soluble and have well-dispersed HSQC spectra at the same protein concentration in the same buffer conditions. Upon superimposing three HSQC spectra, some significant shifts of HSQC peaks were observed (Fig. 3C and 3D). Detailed analysis revealed: 1) CSD values of HSQC peaks upon dissection of hnRNPA1 RRM1-RRM2 (with the largest < 0.3) are much less than those (with the largest close to 1.2 ppm) upon dissection of TDP-43 RRM1-RRM2 (Fig. 1D). 2) residues with large shifts are located on RRM1, linker and RRM2 (Fig. 3B). This observation suggests that in the tethered form, two RRM domains of hnRNPA1 also have some dynamic inter-domain interactions to some degree.

### Thermal stability and ATP-binding of the tethered and isolated hnRNPA1 RRM domains

We further characterized the thermal stability and ATP binding of the tethered and isolated RRM domains of hnRNPA1. The tethered RRM1-RRM2 has only one thermal unfolding transition with Tm of 55 °C, which increased to 58 °C with addition of ATP (Fig. 4A). Intriguingly, the isolated RRM1 without ATP has two thermal unfolding transitions with Tm of 51 and 57 °C respectively. With addition of ATP up to 15 mM, the transition at 51 °C disappeared and only the transition at 57 °C retained. One the other hand, the isolated RRM2 has one thermal unfolding transition with Tm of 60 °C, which increased to 63 °C with the addition of ATP. The results together suggest that unlike TDP-43, the tethering of hnRNPA1 RRM1 and RRM2 domain not only led to no significant destabilization, but in fact resulted in the disappearance of one unfolding transition of the isolated RRM1 at 51 °C, implying that the inter-domain interaction might slightly stabilize the first RRM domain.

**Fig. 4.**
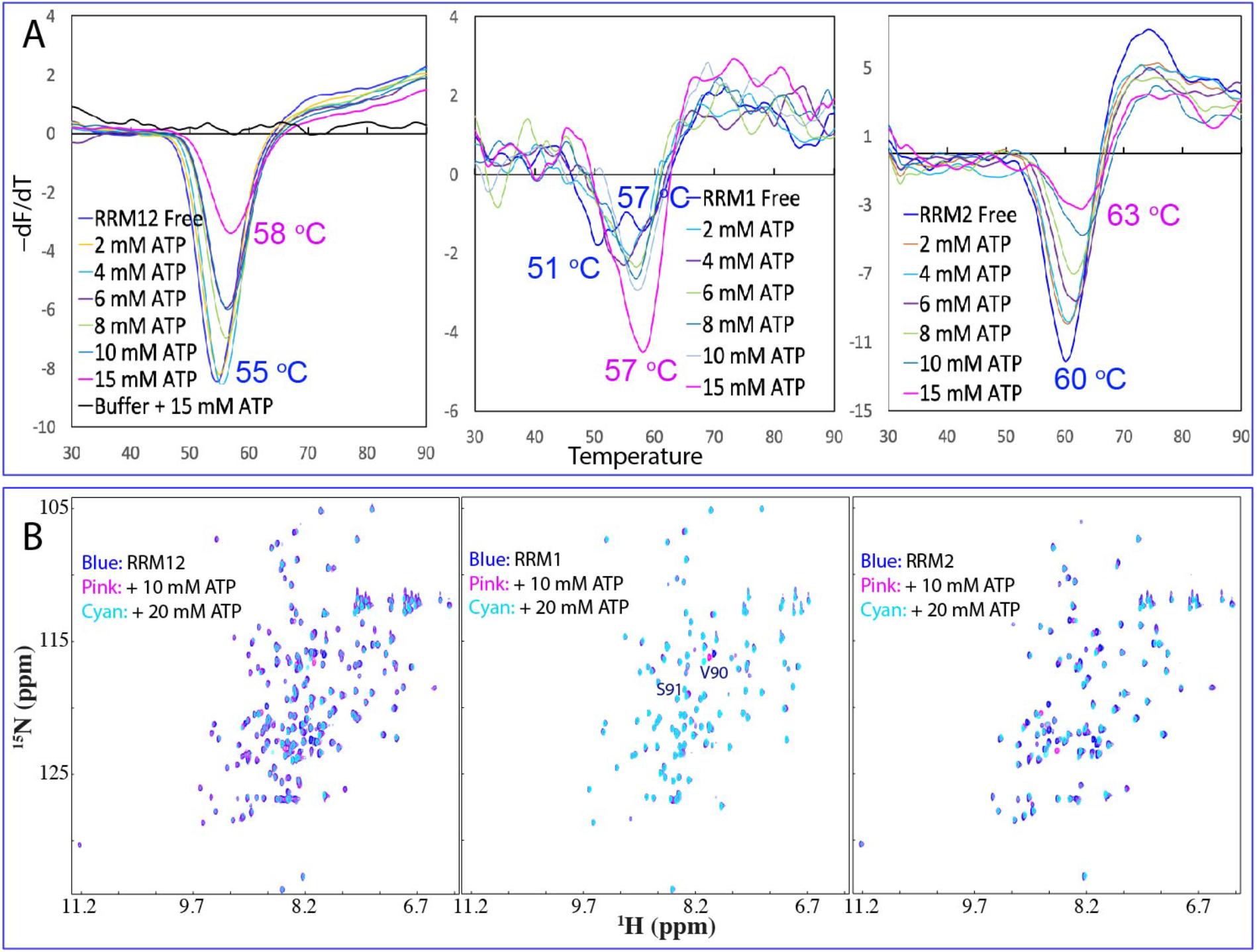
Thermal stability and ATP binding of the hnRNPA1 RRM domains. (A) DSF melting curves of thermal unfolding of the tethered RRM1-RRM2, as well as isolated RRM1 and RRM2 domains in the presence of ATP at different concentrations. (B) ^1^H-^15^N NMR HSQC spectra of the ^15^N-labeled tethered RRM1-RRM2, as well as isolated RRM1 and RRM2 domains at 50 μM (blue) in 10 mM sodium phosphate buffer (pH 6.8) containing 150 mM NaCl and 10 mM DTT in the absence (blue), and in the presence of ATP at 10 (pink) and 20 (cyan) mM.

We further characterized the binding of ATP to the tethered and isolated RRM domains. As shown in Fig. 4B, ATP induced large shifts of many HSQC peaks of the tethered RRM1-RRM2, and detailed analysis revealed that the residues with significant shifts are located on the RRM2 domain except for Arg88, Val90 and Ser91 within RRM1 (Fig. 5A). This was further confirmed by the ATP titrations on the isolated RRM1 and RRM2 domains (Fig. 4B). ATP even with concentrations up to 20 mM only triggered large shifts of two residues Val90 and Ser91 of RRM1 but induced significant shifts of a large set of peaks of the isolated RRM2 domain. Furthermore, the overall patterns of shifted residues of RRM2 are very similar in both tethered and isolated forms (Fig. 5A and 5B).

**Fig. 5.**
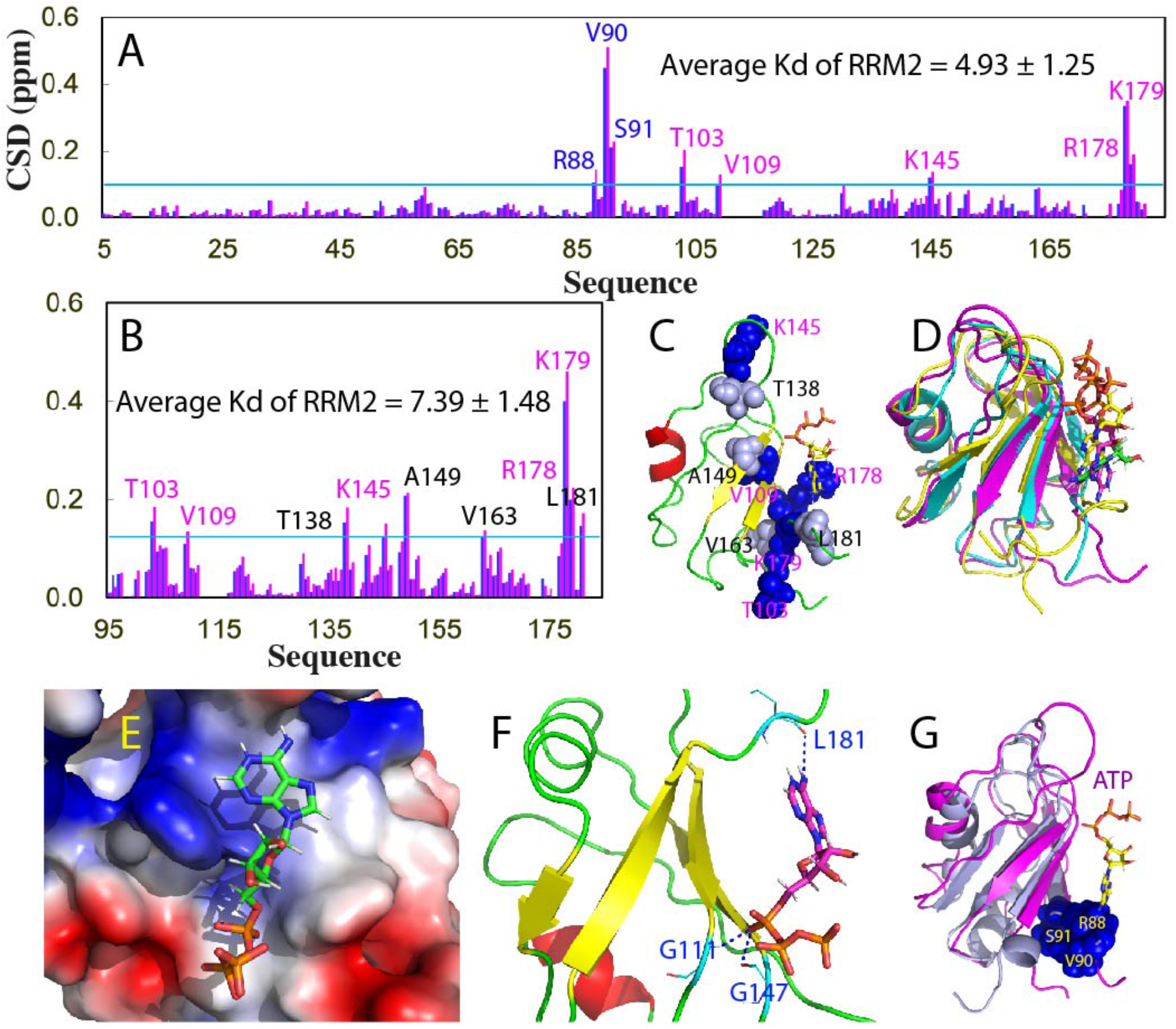
ATP binding of the tethered RRM1-RRM2, as well as isolated RRM2 of hnRNPA1. (A) Residue-specific chemical shift difference (CSD) of the tethered hnRNPA1 RRM1-RRM2 in the presence of ATP at 10 mM (blue) and 20 mM (pink). Significantly shifted residues are labeled, which are defined as those with the CSD values at 20 mM ATP > 0.1 (average value + one standard deviation) (cyan line). (B) Residue-specific chemical shift difference (CSD) of the isolated RRM2 in the presence of ATP at 10 mM (blue) and 20 mM (pink). Significantly shifted residues are labeled, which are defined as those with the CSD values at 20 mM ATP > 0.125 (average value + one standard deviation) (cyan line). The shifted residues identical to those in the tethered form are labeled in pink while the additionally shifted residues are in black. (C) The lowest energy docking model of the ATP-RRM2 complex. The structure of hnRNPA1 RRM2 is displayed in ribbon, while ATP is in sticks. The nine residues with significant CSD values are displayed in spheres and labeled. (D) Superimposition of three complexes of ATP: with hnRNPA1 RRM2 (pink), TDP-43 RRM1 (yellow) and TDP-43 RRM2 (cyan). (E) The ATP-RRM2 complex of hnRNPA1 with the RRM2 structure displayed in the electrostatic potential surface and ATP in sticks. (F) The ATP-RRM complex of hnRNPA1 showing three hydrogen bonds (in blue dotted lines) between ATP and RRM2 atoms. (G) Superimposition of the hnRNPA1 RRM1 and ATP-RRM2 complex with the three significantly shifted RRM1 residue (Arg88, Val90 and Ser91) displayed in spheres.

With the same method we previously used to characterize the ATP-binding to the FUS and TDP-43 RRM domains (20-22,27), we determined Kd value of the ATP binding to hnRNPA1 RRM2 to be 4.93 and 7.39 mM respectively for the tethered and isolated forms (Fig. 5A and 5B). The values are very similar to those for FUS RRM, TDP-43 RRM1 as well as for a non-canonical helix-only RNA-binding domain (28) of hnRNP Q (Kd of 3.14 mM). Interestingly, as observed on TDP-43 RRM domains, the ATP binding affinity to the RRM domains of hnRNPA1 is also higher for the tethered form than for the isolated form. Intriguingly, here ATP is capable of stabilizing the RRM1 domain even without significant binding.

### Visualization of the ATP-RRM2 complex of hnRNPA1

Because of the extremely low binding affinity with Kd of ∼ mM, it is impossible to determine the three-dimensional structure of the ATP-RRM2 complexes by the classic methods of NMR spectroscopy or X-ray crystallography. So here to visualize the complex structure, we utilized the NMR-binding derived constraints to guide the molecular docking with the well-established HADDOCK program (29), as we extensively conducted before on the non-classic ATP-protein complexes (20-22,27).

Fig. 5C presents the lowest-energy docking structure of the ATP-RRM2 complex of hnRNPA1. Overall, this structure is very similar to those of the ATP-RRM1 and ATP-RRM2 complexes of TDP-43 in which ATP occupies a pocket within the conserved surfaces of RRM domains for binding various nucleic acids (Fig. 5D). A close examination reveals that in the ATP-RRM2 complex of hnRNPA1, the aromatic purine ring of ATP has close contacts with the positively-charged surface constituted by the side chains of both RRM2 Arg178 and Lys179 likely to establish *π*-cation interactions on the one hand, as well as with Phe108 and Phe148 likely by establishing *π*-*π* interactions on the other hand (Fig. 5E). Furthermore, the NH of the purine ring of ATP forms a hydrogen bond with the backbone oxygen of Leu181, while the α- and β-phosphate oxyanions of ATP form other two hydrogen bonds respectively with the backbone nitrogen atoms of Gly114 and Gly147 (Fig. 5F).

Although ATP has no complete binding pocket on the RRM1 domain of hnRNPA1, the three residues with large shifts of HSQC peaks induced by adding ATP are also located within the conserved surfaces for RRM domains to bind nucleic acids (Fig. 5G).

### Dynamic behaviours of the TDP-43 tethered and isolated RRM domains

To understand the dynamic basis underlying the coupling of the tethered RRM1-RRM2 of TDP-43, we conducted molecular dynamics (MD) simulations for the tethered as well as isolated RRM1 and RRM2 of TDP-43 with three parallel 50-ns simulations for each constructs. Molecular dynamics simulation is a powerful tool which can not only provide insights into the conformational dynamics that underlies protein functions, but also detect long-range inter-domain correlation motions (30-32) as we previously showed on other proteins (33-36).

I of Fig. 6A presents the root-mean-square deviations (RMSD) of the Cα atoms averaged over three trajectories for the tethered RRM1-RRM2 (black), as well as isolated RRM1 (blue) and RRM2 (pink). The tethered RRM1-RRM2 of TDP-43 has larger RMSD value (4.32 ± 0.39 Å) than those of the isolated RRM1 (3.12 ± 0.35 Å) and RRM2 (2.72 ± 0.32 Å). Furthermore, even with the exclusion of the unstructured N- and C-termini for calculation, RRM1 (104-179) in the tethered form still has larger RMSD value (4.39 ± 0.41 Å) than RRM1 (104-179) in the isolated form (2.51 ± 0.23 Å) (II of Fig. 6A). Similarly, RRM2 (191-261) in the tethered form also has larger RMSD value (3.48 ± 0.37 Å) than RRM2 (104-179) in isolated form (1.74 ± 0.14 Å) (III of Fig. 6A).

**Fig. 6.**
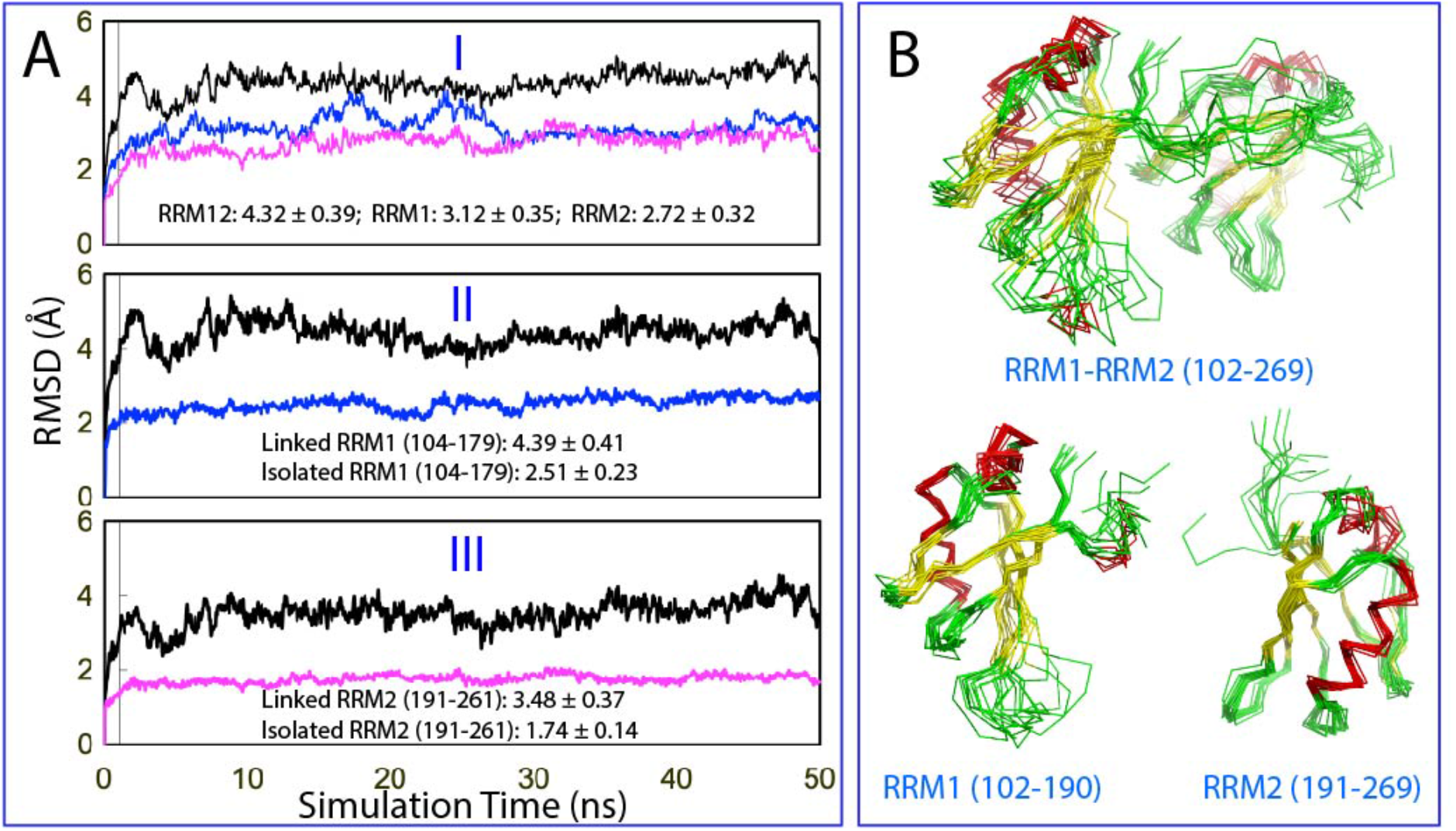
Overall dynamic behaviors of the TDP-43 RRM domains. (A) Average root-mean-square deviations (RMSD) of the Cα atoms over three independent 50-ns MD simulations for the tethered RRM1-RRM2 (black), as well as isolated RRM1 (blue) and RRM2 (pink) (I). for the RRM1 in the tethered form (black) and isolated form (blue) with the unstructured N- and C-termini excluded for calculation (II); as well as for the RRM2 in the tethered form (black) and isolated form (pink) with the unstructured N- and C-termini excluded for calculation (III). (B) Structure snapshots in the first MD simulation with one structure for each 5 ns.

Fig. 6B presents the structure snapshots in the first MD simulations for the tethered RRM1-RRM2 as well as isolated RRM1 and RRM2, clearly indicating that the structures of the tethered RRM1-RRM2 are more fluctuating than those of the isolated RRM1 or RRM2, completely consistent with the RMSD results. Similar dynamic behaviours are also reflected by the root-mean-square fluctuations (RMSF) of the Cα atoms averaged over three trajectories (Fig. 7). As shown in Fig. 7A, while the residue-specific fluctuations of RRM1 are very similar in both tethered and isolated forms, those of RRM2 in the tethered form are higher than those in the isolated forms. As a consequence, the majority of the residues with significant differences of RMSF between tethered and isolated forms are located in RRM2 (Fig. 7B and 7C), which is in general consistent with the dissection-induced effects as detected by NMR (Fig. 1D).

**Fig. 7.**
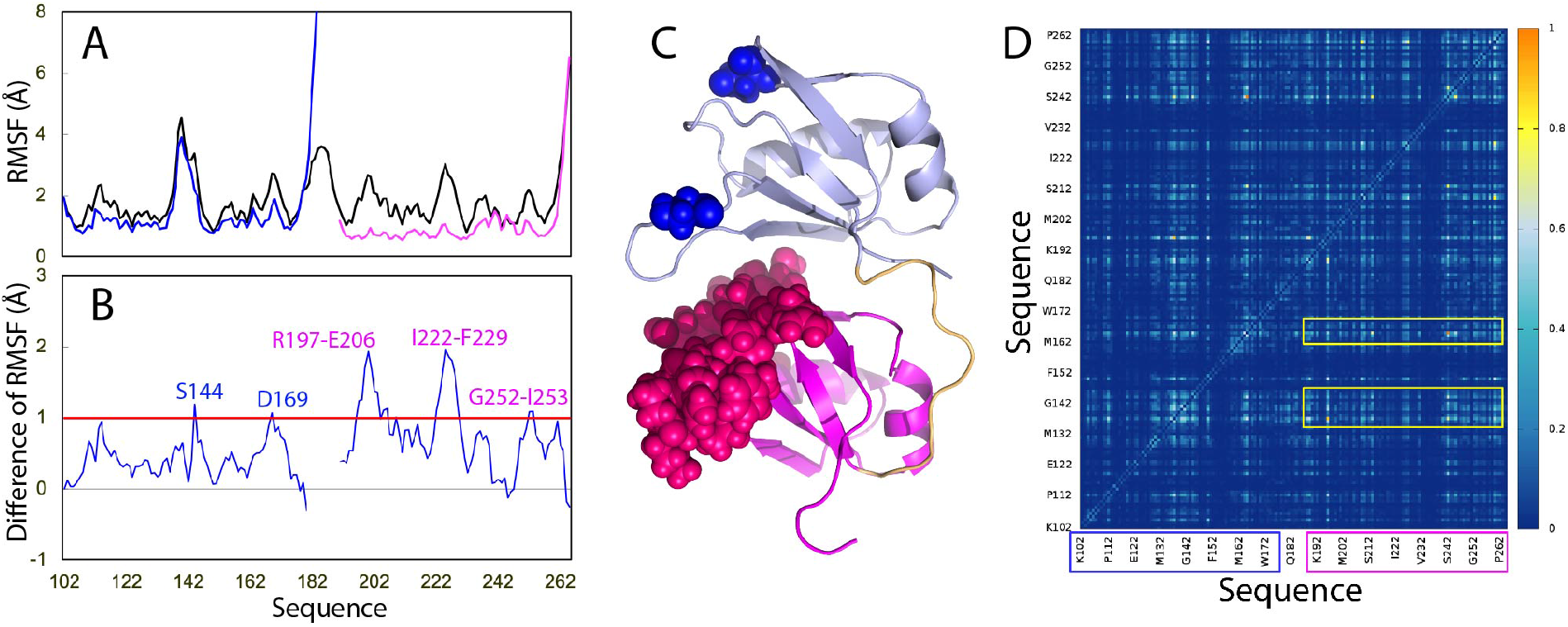
Residue-specific dynamic behaviors of the TDP-43 RRM domains. (A) Averaged root-mean-square fluctuations (RMSF) of the Cα atoms computed over three independent MD simulations for the tethered RRM1-RRM2 (black), as well as isolated RRM1 (blue) and RRM2 (pink). (B) The difference of the average RMSF between the tethered RRM1-RRM2, and isolated RRM1/RRM2. The residues are labeled for those with significant difference of RMSF (> 1.0; average + one STD). (C) Structure of the TDP-43 RRM1-RRM2 with the residues displayed in spheres which show significant difference of RMSF between the tethered and isolated forms. (D) Mutual information matrix calculated from three parallel MD simulation data of the tethered TDP-43 RRM1-RRM2 by MutInf, with residues having significant inter-domain correlation motions highlighted by yellow boxes. The RRM1 sequence is indicated by blue box and RRM2 sequence in pink box.

MutInf represents an entropy-based approach to analyze ensembles of protein conformers, such as those from molecular dynamics simulations by using internal coordinates and focusing on dihedral angles. In particular, this approach is particularly applicable for those in which conformational changes are subtle (33). Briefly, this approach utilizes second-order terms from the configurational entropy expansion, called the mutual information, to identify pairs of residues with correlated conformations, or correlated motions. Fig. 7D shows the normalized correlation motion matrix of the tethered RRM1-RRM2 of TDP-43. Interestingly, the correlation motions exist not only within RRM1 or RRM2 domain, but also between two domains. In particular, the RRM1 residues around Gly142 and Met162 have extensive correlation motions with many RRM2 residues. Furthermore, the strength of the inter-domain correlation motions has no significant difference from that of the intra-domain motions, suggesting that the tethered RRM1-RRM2 indeed behaves as a coupled dynamic unit. This rationalizes the DSF results that the tethered RRM1-RRM2 of TDP-43 only has one thermal unfolding transition although the isolated RRM1 and RRM2 have their own transitions with very different Tm values.

### Dynamic behaviours of the hnRNPA1 tethered and isolated RRM domains

We also conducted 50-ns molecular dynamics (MD) simulations for the tethered as well as isolated RRM1 and RRM2 of hnRNPA1 with three parallel simulations for each constructs. I of Fig. 8A presents the root-mean-square deviations (RMSD) of the Cα atoms averaged over the three trajectories for the tethered RRM1-RRM2 (black), as well as isolated RRM1 (blue) and RRM2 (pink). Interestingly, the tethered RRM1-RRM2 of hnRNPA1 also has larger RMSD value (4.87 ± 1.05 Å) than those of RRM1 (3.39 ± 0.48 Å) and RRM2 (3.66 ± 0.49 Å). Furthermore, although the unstructured N- and C-termini were not included for calculation, the RRM1 (9-91) in the tethered form still has larger RMSD value (4.69 ± 1.23 Å) than RRM1 (9-91) in the isolated form (1.97 ± 0.21 Å) (II of Fig. 8A). Similarly, RRM2 (105-180) in the tethered form also has larger RMSD value (4.25 ± 0.92 Å) than RRM2 (105-180) in the isolated form (1.77 ± 0.19 Å) (III of Fig. 8A).

**Fig. 8.**
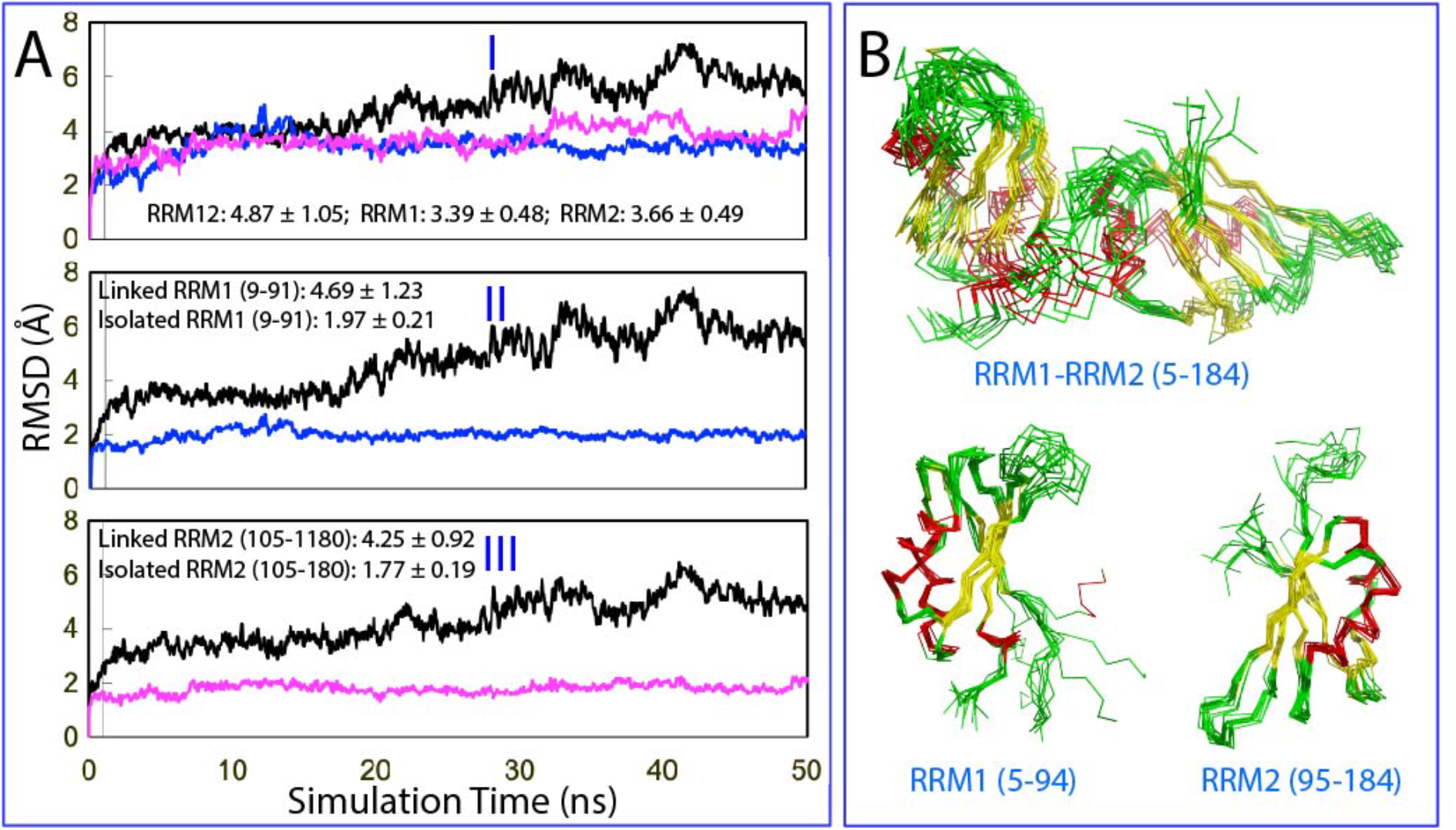
Overall dynamic behaviors of the hnRNPA1 RRM domains. (A) Average root-mean-square deviations (RMSD) of the Cα atoms over three independent 50-ns MD simulations for the tethered RRM1-RRM2 (black), as well as isolated RRM1 (blue) and RRM2 (pink) (I). for the RRM1 in the tethered form (black) and isolated form (blue) with the unstructured N- and C-termini excluded for calculation (II); as well as for the RRM2 in the tethered form (black) and isolated form (pink) with the unstructured N- and C-termini excluded for calculation (III). (B) Structure snapshots in the first MD simulation with one structure for each 5 ns.

Fig. 8B presents the structure snapshots in the first MD simulations for the tethered RRM1-RRM2 as well as isolated RRM1 and RRM2 of hnRNPA1, showing that the structures of the tethered RRM1-RRM2 are indeed more fluctuating than those of the isolated RRM1 and RRM2, completely consistent with the RMSD results. Noticeably, different from those observed for the TDP-43 RRM domains (Fig. 7), the residue-specific RMSF values of both RRM1 and RRM2 of hnRNPA1 in the tethered forms are larger than those of isolated RRM1 and RRM2 (Fig. 9A). Consequently, the residues with significant differences of RMSF between the tethered and isolated forms are located on both RRM1 and RRM2 (Fig. 9B and 9C), which is in general consistent with the dissection-induced perturbation as detected by NMR (Fig. 3D).

**Fig. 9.**
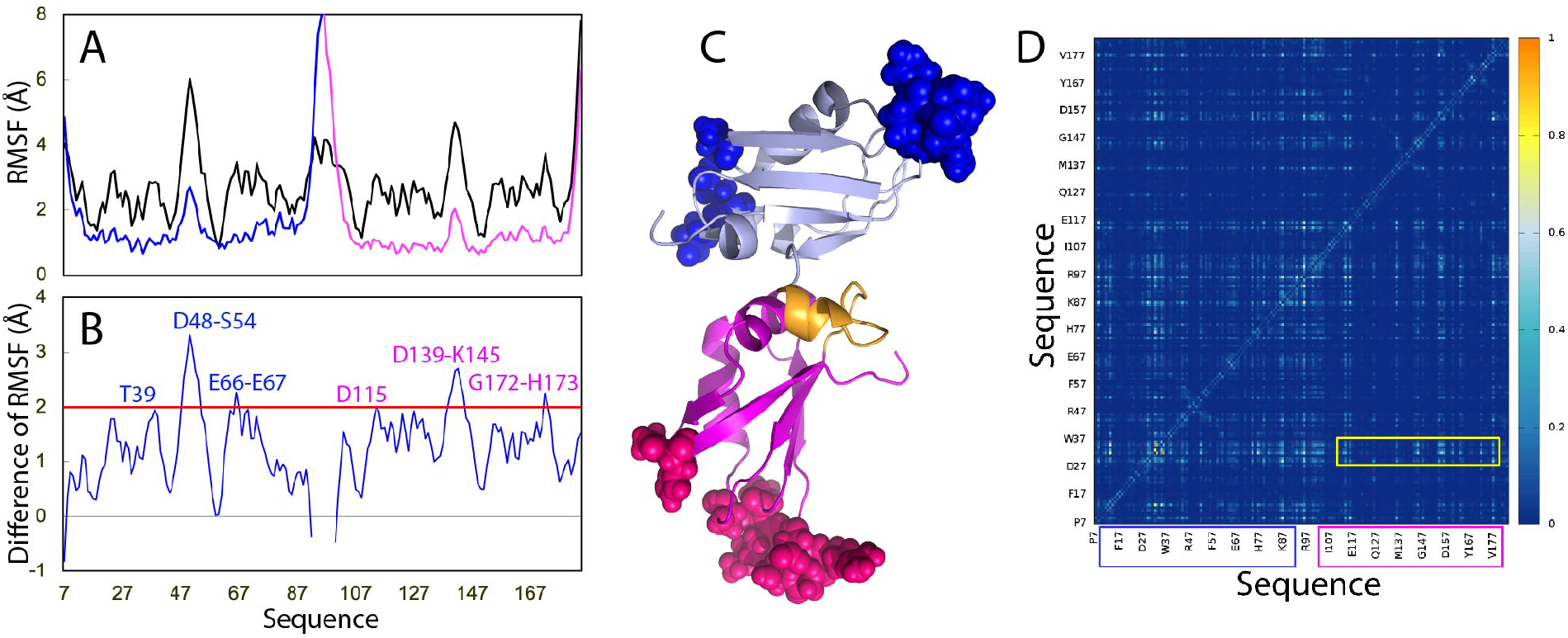
Residue-specific dynamic behaviors of the hnRNPA1 RRM domains. (A) Averaged root-mean-square fluctuations (RMSF) of the Cα atoms computed over three independent MD simulations for the tethered RRM1-RRM2 (black), as well as isolated RRM1 (blue) and RRM2 (pink). (B) The difference of the average RMSF between the tethered RRM1-RRM2, and isolated RRM1/RRM2. The residues are labeled for those with significant difference of RMSF (> 2.0; average + one STD). (C) Structure of the hnRNPA1 RRM1-RRM2 with the residues displayed in spheres which show significant difference of RMSF between the tethered and isolated states. (D) Mutual information matrix calculated from three parallel MD simulation data of the tethered hnRNPA1 RRM1-RRM2 by MutInf, with residues having significant inter-domain correlation motions highlighted by yellow box. The RRM1 sequence is indicated by blue box and RRM2 sequence in pink box.

Fig. 9D presents the normalized correlation motion matrix of the tethered RRM1-RRM2 of hnRNPA1. Although the correlation motions still exist between two domains, the inter-domain correlation motions are mainly between the RRM1 residues Arg31-Ser32 and RRM2 residues. The strength of the correlation motions for the hnRNPA1 RRM1-RRM2 is weaker than that observed for the TDP-43 RRM1-RRM2 (Fig. 7D). This suggests that the coupling of hnRNPA1 RRM1 and RRM2 domains might be weaker than that for TDP-43 RRM1 and RRM2 domains.

## Discussion

Protein aggregation/fibrillation has been now established to be the universal hallmark of an increasing spectrum of human diseases beyond neurodegenerative diseases, which also include cardiac dysfunction, eye cataract, degeneration of muscle and bone, as well as aging down to *E. coli* (24,37-48). Out of various factors that modulate aggregation/fibrillation of the folded proteins, two key determinants are thermodynamic stability and conformational dynamics. In human genome, many RRM-containing proteins such as TDP-43 and hnRNPA1 have two tethered RRM domains, which have been demonstrated to also play a key role in the disease-causing aggregation/fibrillation. Although the observations imply that the tethered TDP-43 RRM domains are particularly prone to aggregation/fibrillation, so far there has been no systematic study to understand the underlying mechanisms, which, however, are of both fundamental and therapeutic interest.

In the present study, by both experimental and computational approaches we conducted the first systematic study to characterize the thermal stability and ATP-binding by DSF and NMR, followed by MD simulations to assess the conformational dynamics of two RRM domains of TDP-43 and hnRNPA1 in both tethered and isolated forms. Very unexpectedly, the results showed that the isolated TDP-43 RRM1 and RRM2 domains have Tm of 57 and 59 °C respectively, while the tethered form has only one unfolding transition with Tm significantly reduced to only 49 °C. This set of results clearly indicates that the tethering induced the significant coupling of two RRM domains of TDP-43, as well as dramatic destabilization. By contrast, no significant destabilization was observed for the tethering of two RRM domains of hnRNPA1. The results thus underscore the extreme complexity of the tethering-induced effects even for the tandem RRM domains of the different members within the same hnRNP protein family. In a general context, the dramatic destabilization for the tethered TDP-43 RRM domains is also very unusual because previously the tethering was demonstrated to have either no significant effect or even to stabilize the tethered domains. Only recently it was found that the tethering may also destabilize the protein domains (49-51), as exemplified by the ubiquitination-induced destabilization of the modified proteins, which was proposed to function to facilitate their degradation (52). Interestingly, the tethering-induced destabilization has been proposed to evolve from the trade-offs for functions (51). In this regard, it is of great interest in the future to define the functional role of the unique destabilization for tandem TDP-43 RRM domains. Nevertheless, this unexpected destabilization certainly contributes to the unusually high tendency of TDP-43 in aggregation/fibrillation, which has been well established to lead to a variety of human diseases by “loss of function” or/and “gain of function”.

The tethering-induced effects for TDP-43 and hnRNPA1 RRM domains appear to result mainly from their inter-domain interactions. Residue-specific NMR data showed that the dissection-induced perturbation for TDP-43 RRM domains is much more profound than that for hnRNPA1 RRM domains, while MD simulations further revealed that the inter-domain correlation motions of TDP-43 RRM domains are more extensive than those for hnRNPA1 RRM domains. Therefore, experimental and simulation results together provide a mechanism to rationalize the observation that the tethered TDP-43 RRM domains could become highly coupled with only one unfolding transition with Tm much lower than those of their isolated ones. On the other hand, it remains an extreme challenge in the future to understand why the inter-domain interactions significantly destabilize the two RRM domains of TDP-43 but lead to even a slight stabilizing effect on the RRM1 domain of hnRNPA1.

Mysteriously, all cells maintain very high ATP concentrations of 2-12 mM, much higher than those required for its previously-known functions (52-56), although the majority of ATP needs to be produced by very complex supramolecular machineries embedded in membranes (53). Only recently, it was decoded that ATP with concentrations > 5 mM acts to hydrotropically dissolve liquid-liquid phase separation (LLPS), aggregation/fibrillation (54), which appears to operate at a proteome-wide scale (55). We further found that by weak but specific binding with Kd of ∼mM, ATP is also able to modulate LLPS of intrinsically disordered domains in the two-stage manner (56,57). Remarkably, ATP can inhibit amyloid fibrillation not only for the tethered TDP-43 RRM1-RRM2 by enhancing its thermal stability (21), but also for the single FUS RRM domain by blocking the dynamic opening of the RRM fold without alternation of its thermal stability (20). Here, again we found that ATP can specifically bind the hnRNPA1 RRM2 domain in both tethered and isolated contexts with the affinity and complex structure very similar to those of the FUS and TDP-43 RRM domains. Moreover ATP can even enhance the thermal stability of the hnRNPA1 RRM1-RRM2 domains. In particular, our previous and current results together suggest that the affinities of the ATP binding to the tethered forms of both TDP-43 and hnRNPA1 RRM domains are higher than those to the isolated forms. This may be mainly due to the higher conformational dynamics of the tethered RRM domains than those of the isolated forms as uncovered by MD simulations, which thus provides ATP the higher dynamic accessibility to the binding pockets in the tethered RRM domains. In light of the previous results with the FUS RRM domain that its high conformational dynamics allow the opening of the RRM fold, thus leading to aggregation/fibrillation but that the ATP binding even without enhancing the stability is sufficient to kinetically block the opening, here the results that ATP can bind the conserved pockets of TDP-43 and hnRNPA1 RRM domains highlight that ATP may play a general role in preventing the pathological aggregation/fibrillation of the RRM-containing proteins containing more than one RRM domain.

In summary, in the present study we showed for the first time that very unexpectedly upon tethering, two TDP-43 RRM domains become highly coupled but dramatically destabilized with the Tm reduction of ∼8 °C. By contrast, no significant destabilization occurs for the tethering of two hnRNPA1 RRM domains. Mechanistically, the tethering-induced effects appear to mainly result from the inter-domain interactions between two RRM domains as reflected by NMR and MD simulation results. Moreover, we showed that ATP can specifically bind TDP-43 and hnRNPA1 RRM domains with the affinity to the tethered forms higher than to the isolated forms. Results together thus suggest that ATP, the universal energy currency, may also play a general role in preventing aggregation/fibrillation of RRM-containing proteins, which has been extensively identified to cause an increasing spectrum of human diseases beyond neurodegenerative diseases. Therefore, our study provides one potential mechanism to rationalize the observation that upon being aged, the risk of protein aggregation-causing diseases increases most likely also because ATP concentrations gradually reduce in all cells during aging (53,54).

## Methods

### Cloning, expression and purification of the tethered and isolated RRM domains of TDP-43 and hnRNPA1

Previously the expression vectors of TDP-43 RRM1-RRM2 (102-269), RRM1 (102-191) and RRM2 (191-269) have been constructed and their expression and purification were established (21,22,58). On the other hand, the expression vector of the tethered RRM1-RRM2 (5-184) of hnRNPA1 were purchased from a local company (Bio Basic Asia Pacific Pte Ltd), which was subsequently dissected into the isolated RRM1 (5-95) and RRM2 (94-184) by PCR with designed primers.

The six recombinant proteins were expressed in *E. coli* BL21 cells with IPTG induction at 20 °C overnight. They were found all in the supernatant, and therefore were purified by a Ni^2+^-affinity column (Novagen) under native conditions. Subsequently the on-gel cleavage by thrombin was conducted and the eluted fractions containing the RRM proteins were further purified by a heparin column to remove nucleic acids followed by FPLC purification with either a Superdex-75 or a Superdex-200 column.

Here we followed our previous protocol to generate isotope-labeled proteins for NMR studies (20-24,58). Briefly, the bacteria were grown in M9 medium with addition of (^15^NH_4_)_2_SO_4_ for ^15^N-labeling. The protein concentrations were determined by the UV spectroscopic method in the presence of 8 M urea, under which the extinct coefficient at 280 nm of a protein can be calculated by adding up the contribution of Trp, Tyr, and Cys residues (59).

ATP was purchased from Sigma-Aldrich with the same catalog numbers as previously reported. MgCl_2_ was added into ATP for stabilization by forming the ATP-Mg complex (20-22,53). The fluorescent dye SYPRO Orange (S5692-50UL) was purchased from Sigma-Aldrich. The protein samples, as well as ATP, were all prepared in 10 mM sodium phosphate buffer containing 10 mM DTT and 150 mM NaCl with a final pH adjusted to 6.8.

### Determination of thermal stability

As we previously showed (20-24), ATP triggered very high non-specific noise in CD spectroscopy and quenched the intrinsic fluorescence of exposed Trp residues, here again we used differential scanning fluorimetry (DSF) as we previously reported to determine the thermodynamic stability of RRM1-RRM2, RRM1 and RRM2 domains of TDP-43 and hnRNPA1 at 10 μM in 10 mM sodium phosphate buffer containing 10 mM DTT and 150 mM NaCl (pH 6.8) with addition of ATP at different concentrations.

DSF experiments were performed using the CFX384 Touch™ Real-Time PCR Detection System from BIO-RAD, following the SYBR green melting protocol to obtain Tm value (21-24). Briefly, in a single well of a 384-well PCR plate, a 10 µl reaction solution was placed, which contains the RRM12, or RRM1 or RRM2 domain at 10 µM, ATP at different concentrations, and 10×SYPRO Orange in 10 mM sodium phosphate buffer containing 150 mM NaCl (pH 6.8). The program in Real-Time PCR instrument was set to be SYBR green and run temperature scan from 30-90 °C with the increment of 1 °C per minute. Upon completion, the obtained thermal unfolding curves were displayed as the first derivatives (dF/dT) by the RT-PCR software Bio-Rad CFX Manager 3.0.

### NMR characterizations of the ATP binding

All NMR experiments were acquired at 25 °C on an 800 MHz Bruker Avance spectrometer equipped with pulse field gradient units and a shielded cryoprobe as we described previously (20-24). For NMR HSQC titration studies of the interactions of RRM1-RRM2, RRM1 and RRM2 with ATP, two dimensional ^1^H-^15^N NMR HSQC spectra were collected on the ^15^N-labelled samples at a protein concentration of 50 µM in 10 mM sodium phosphate buffer containing 10 mM DTT and 150 mM NaCl (pH 6.8) at 25 °C in the absence and in the presence of ATP at concentrations of 0.5, 1.0, 2.0, 4.0, 6.0, 8.0, 10.0, 12.0, 14.0, 16.0, 18.0, and 20.0 mM.

### Calculation of CSD and data fitting to obtain Kd

Sequential assignments of the tethered and isolated RRM1 and RRM2 domains of TDP-43 and hnRNPA1 were achieved based on the previously deposited chemical shifts: the tethered TDP-43 RRM1-RRM2 domain (BMRB ID of 19290 and 27613), the isolated RRM1 (BMRB ID of 18765) and RRM2 (BMRB ID of 19922), as well as the tethered hnRNPA1 RRM1-RRM2 domain (BMRB ID of 18728).

To calculate chemical shift difference (CSD), the HSQC spectra were superimposed for the ^15^N-labeled RRM1-RRM2, RRM1 and RRM2 domains collected in the absence and in the presence of ATP at different concentrations. Subsequently, the shifted HSQC peaks could be identified and further assigned to the corresponding RRM residues based on the sequential assignments. The chemical shift difference (CSD) was calculated by an integrated index calculated by the following formula:

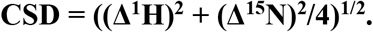

In order to obtain residue-specific dissociation constant (Kd), we fitted the shift traces of the residues with significant shifts of HSQC peaks (CSD > average + STD), by using the one binding site model (20-23,27) with the following formula:

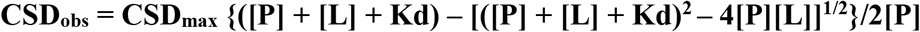

Here, [P] and [L] are molar concentrations of RRM domains and ligands (ATP) respectively.

### Molecular docking

The structure model of the ATP-RRM2 complex of hnRNPA1 was constructed by use of HADDOCK software (20-22, 29), which makes use of CSD data to derive the docking with various degrees of flexibility. Briefly, five residues of hnRNPA1 RRM2 with significant CSD values were set to be active residues and HADDOCK docking procedure for the complexes was performed at three stages: (1) randomization and rigid body docking; (2) semi-flexible simulated annealing; (3) flexible explicit solvent refinement. The ATP-RRM2 structure with the lowest energy score were selected for the detailed analysis and display by Pymol (The PyMOL Molecular Graphics System, Version 0.99rc6 Schrödinger, LLC).

### Molecular dynamics (MD) simulations

For MD simulations, the NMR structures of RRM1-RRM2 of TDP-43 (PDB ID of 4BS2) and hnRNPA1 (PDB ID of 2LYV) were selected as the tethered models while their isolated RRM1 and RRM2 models were obtained by dissecting the two structures into the isolated RRM domains.

The simulation setting was previously reported (34-36). Briefly, the simulation cell is a periodic cubic box with a minimum distance of 10 Å between the protein and the box walls to ensure the protein would not directly interact with its own periodic image given the cutoff. The water molecules, described using the TIP3P model, were filled in the periodic cubic box for the all atom simulation. To neutralize the system, some Na+ and Cl-ions were randomly placed far away from the surface of the protein.

Three independent 50-ns MD simulations were performed for each of six constructs: namely the tethered RRM1-RRM2, isolated RRM1 and RRM2 of TDP-43 as well as of hnRNPA1 by the program GROMACS (60) with the AMBER-03 all-atom force field (61). The long-range electrostatic interactions were treated using the fast particle-mesh Ewald summation method, with the real space cutoff of 9 Å and a cutoff of 14 Å was used for the calculation of van der Waals interactions. The temperature during simulation was kept constant at 300 K by Berendsen’s coupling. The pressure was held at 1 bar. The isothermal compressibility was 4.5*10−5 bar-1. The time step was set as 2 fs. Prior to MD simulations, all the initial structures were relaxed by 1000 steps of energy minimization using steepest descent algorithm, followed by 100 ps equilibration with a harmonic restraint potential applied to all the heavy atoms of the proteins.

### Analysis of correlation motions

To detect the coupling of the RRM1 and RRM2 domains, we calculated the correlation matrixes of three parallel simulations of the tethered RRM1-RRM2 of both TDP-43 and hnRNPA1 by a recently established approach called MutInf (32). Briefly, MutInf represents an entropy-based approach to analyze ensembles of protein conformers, such as those from molecular dynamics simulations by using internal coordinates and focusing on dihedral angles. In particular, this approach is even applicable in cases for which conformational changes are subtle. Briefly, this approach utilizes second-order terms from the configurational entropy expansion, called the mutual information, to identify pairs of residues with correlated conformations, or correlated motions, in an equilibrium ensemble. In the present study, the normalized matrix values were used, and 0.3 was set up to be the threshold value to determine the pairs of highly correlated residues.

## Acknowledgements

This work was supported by Ministry of Education (MOE), Singapore Tier 1 Grant R-154-000-B92-114.

## Author contributions

J.S. and M.D. conceived and designed the experiments. M.D. Y.L. and J.S. performed the research, analyzed the data. J.S. wrote the manuscript.

## Notes

### Competing Interest Statement

The authors have declared no competing interest.

